# ‘Normal’ hearing thresholds and fundamental auditory grouping processes predict difficulties with speech-in-noise perception

**DOI:** 10.1101/578393

**Authors:** Emma Holmes, Timothy D. Griffiths

## Abstract

Understanding speech when background noise is present is a critical everyday task that varies widely among people. A key challenge is to understand why some people struggle with speech-in-noise perception, despite having clinically normal hearing. Here, we developed new figure-ground tests that require participants to extract a coherent tone pattern from a stochastic background of tones. These tests dissociated variability in speech-in-noise perception related to mechanisms for detecting static (same-frequency) patterns and those for tracking patterns that change frequency over time. In addition, elevated hearing thresholds that are widely considered to be ‘normal’ explained significant variance in speech-in-noise perception, independent of figure-ground perception. Overall, our results demonstrate that successful speech-in-noise perception is related to audiometric thresholds, fundamental grouping of static acoustic patterns, and tracking of acoustic sources that change in frequency. Crucially, measuring both peripheral (audiometric thresholds) and central (grouping) processes is required to adequately assess speech-in-noise deficits.

## Introduction

From ordering a coffee in a busy café to maintaining a conversation while walking down the street, we are often required to communicate when background noise is present (“speech-in-noise perception”). These situations are known to be challenging for people with hearing impairment (Marrone et al., 2008). Yet, it has been estimated that 5–15% (Cooper and Gates, 1991; Hind et al., 2011; Kumar et al., 2007) of people who seek clinical help for their hearing have normal hearing thresholds; the main problem they report is difficulty understanding speech in noisy places. Why some people struggle with speech-in-noise perception, despite displaying no clinical signatures of peripheral hearing loss, has been difficult to elucidate.

One idea that has gained recent attention is that speech-in-noise difficulty is related to cochlear synaptopathy (Kujawa and Liberman, 2009): damage to the synapses between the cochlea and auditory nerve fibres. Cochlear synaptopathy primarily affects high-threshold, low-spontaneous-rate fibres (Furman et al., 2013), which are thought to be important for supra-threshold perception, providing a mechanism by which speech perception could be distorted without elevating hearing thresholds (Oxenham, 2016). In animal models, cochlear synaptopathy has been linked to changes in electrophysiological measures, such as the auditory brainstem response (ABR; Hickox et al., 2017; Shaheen et al., 2015) and envelope following response (EFR; Shaheen et al., 2015). However, in humans, links between cochlear synaptopathy and electrophysiological measures have not been established (Hickox et al., 2017) and proposed measures of cochlear synaptopathy do not correlate well (Guest et al., 2019). Some studies relate poorer speech-in-noise performance to lower EFR (Bharadwaj et al., 2015; Ruggles et al., 2011) or lower ABR wave 1 (Bramhall et al., 2015) amplitudes, but others have found no evidence for an association with EFRs (Guest et al., 2018) or ABRs (Fulbright et al., 2017; Guest et al., 2018). These mixed results imply that either cochlear synaptopathy is not a prominent source of variability in speech-in-noise perception in humans, or we do not currently have a good way to assess it (see Guest et al., 2018).

Another possibility, which has been explored in less detail, is that difficulty with speech-in-noise perception originates from impaired central auditory processes. To understand speech when other conversations are present, we must successfully group parts of the acoustic signal that belong to target speech, sustain our attention on these elements, and hold relevant speech segments in working memory. Some previous studies have linked speech-in-noise perception to working memory and attention, although no single test produces reliable correlations (see Akeroyd, 2008). Furthermore, experiments that have provided evidence for this relationship have typically used cognitive test batteries developed for clinical application, and these are non-specific: impaired performance could be attributed to a variety of cognitive and perceptual mechanisms (Crowe, 1998). Thus, we do not fully understand the extent to which central factors contribute to variability in speech-in-noise perception among people with normal hearing, or which factors are important for explaining between-subject variability.

Here, we hypothesised that individual variability in speech-in-noise perception might be related to central auditory processes for grouping sound elements as belonging to target or competing speech. To this aim, we investigated whether specific tests of auditory figure-ground perception predict speech-in-noise performance. Our figure-ground tests were based on an established test that assesses the ability to detect pure tones that are fixed in frequency across time (the ‘figure’) among a ‘background’ of random frequency tones (Teki et al., 2013, 2011; O’Sullivan et al., 2015). In the prototype stimulus, the figure and background components are acoustically identical at each time window and cannot be distinguished; successful figure detection requires the listener to group tones over time, which could be accomplished by a temporal coherence mechanism (Teki et al., 2013). Improved detectability of these figures (by increasing their coherence or duration) has been associated with increased activity in the superior temporal sulcus, inferior parietal sulcus, and the right planum temporale (Teki et al., 2011; Tóth et al., 2016). We predicted that people who are worse at figure-ground perception are also worse at speech-in-noise perception.

Our new tests differed from previous figure-ground tasks, which asked participants to detect whether or not a figure was present among a ‘background’ of random-frequency tones. Instead, we presented a two-interval forced choice (Fechner, 1860) discrimination task, in which a 300-ms ‘gap’ occurred in the figure or background components. Both figure and background were present in all stimuli, and participants heard two stimuli, presented sequentially, on each trial. They were asked to discriminate which stimulus (first or second interval) contained a gap in the figure components. Successful performance in this task requires participants to successfully extract the figure from background components and cannot be performed based on global stimulus characteristics. Performance in these tasks is unlikely to be related to gap discrimination thresholds, which are of an order of magnitude lower than the gap duration we used here (Shailer and Moore, 1983). Instead, performing the task requires listeners to determine whether the gap occurred in the figure or background components, which will be more difficult if the figure components are more difficult to group. This task is closer to speech-in-noise perception than the figure detection task used in previous studies, yet isolates grouping processes from semantic and other cognitive demands required for speech-in-noise perception.

To tease apart different grouping mechanisms, we tested three classes of ‘figure’, in which the relationship between frequency elements differed (see Figure 1). Similar to classic figure-ground stimuli, one class of figure contained three components that remained the same frequency over time. In a second class, the three components were multiples of the first formant extracted from naturally spoken sentences. These figures changed frequency over time, similar to the first formant of natural speech, and all figure components changed frequency at the same rate. The third class of figure was constructed from the first, second, and third formants extracted from the same spoken sentences: the three figure components changed frequency at different rates, similar to the formants in speech. For all stimuli, the figure and background were constructed from 50-ms tones, rendering the formants unintelligible, and ensuring that the three figure classes had similar acoustic properties.

**Figure 1.**
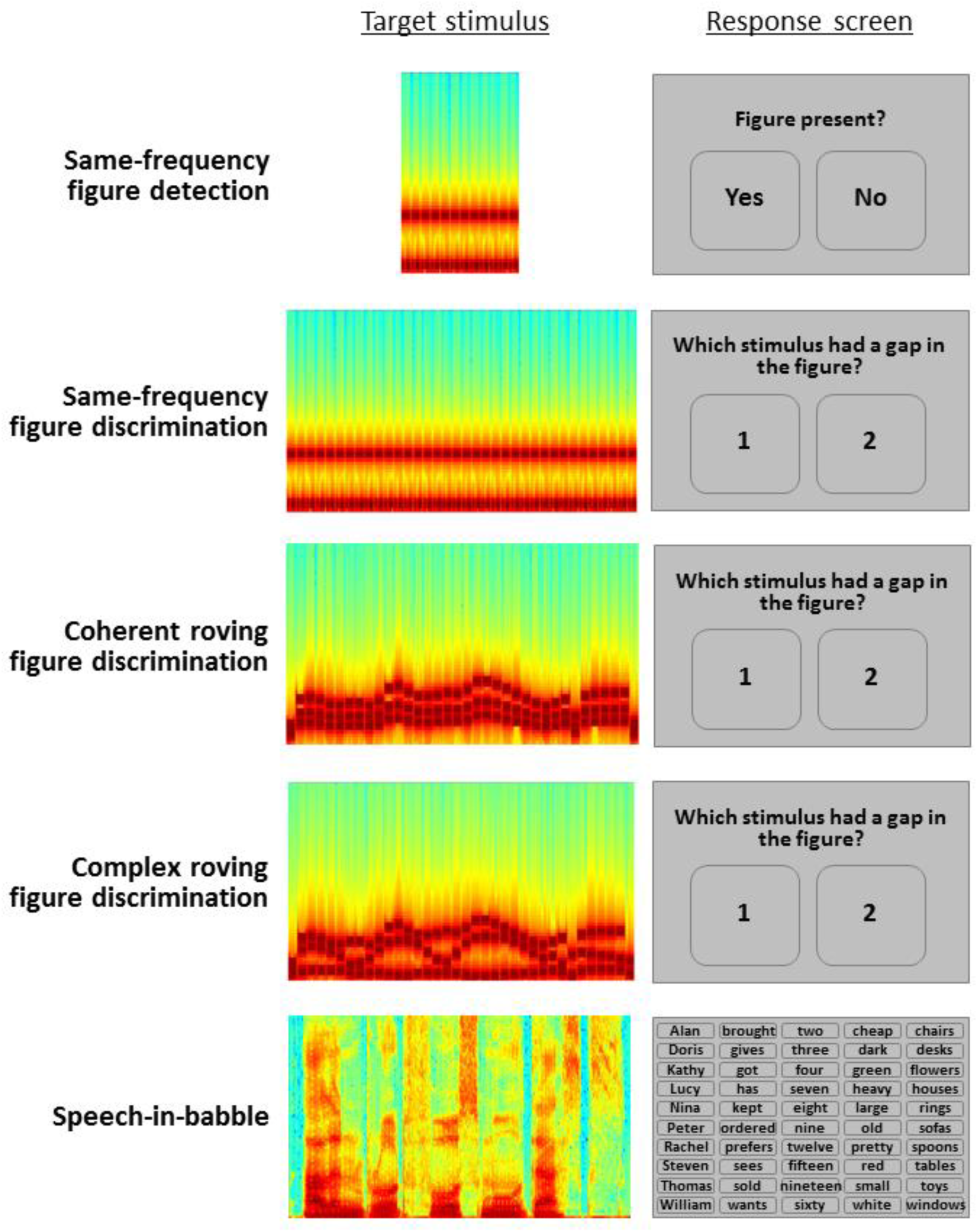
Schematic of design. Left panel: Example spectrogram of one target stimulus for each task (figure or speech). Right panel: Schematic of response screen for each task.

We recruited participants who had normal hearing and excluded any who would be considered to have mild hearing loss (defined as six-frequency averages at 0.25–8 kHz ≥ 20 dB HL in either ear). Despite this, we also sought to examine whether sub-clinical variability in audiometric thresholds, which index aspects of peripheral processing, contain information relevant for predicting speech-in-noise perception. This follows from the idea that speech-in-noise difficulties may arise from changes to the auditory periphery that are related to, but which precede, clinically relevant changes in thresholds (see Pienkowski, 2016). Given that high-frequency thresholds are suspected to deteriorate first in age-related hearing loss (Wiley et al., 2008), we tested thresholds at 4–8 kHz.

Each participant performed a speech-in-noise task, figure-ground tasks, and audiometric testing. Our results demonstrate that clinically ‘normal’ hearing thresholds contain useful information for predicting speech-in-noise perception. In addition, performance on our new figure-ground tasks explains significant variability in speech-in-noise performance that is not explained by audiometric thresholds. Overall, the results demonstrate that different people find speech-in-noise perception difficult for different reasons: the limitation arises for some participants due to slightly elevated thresholds for detecting quiet sounds, for others due to central processes related to the fundamental grouping of fixed-frequency acoustic patterns, and for others due to difficulty tracking objects that change frequency over time.

## Results

Figure 2A illustrates the correlations between performance on the speech-in-babble task and audiometric thresholds and between performance on the speech-in-babble and figure-ground tasks. The shaded region at the top of the graph displays the noise ceiling, defined as the correlation between thresholds measured in two separate blocks of the speech-in-babble task (*r* = .69, *p* < .001; 95% CI = .56–.78). For all subsequent analyses, we averaged thresholds across these two blocks.

**Figure 2.**
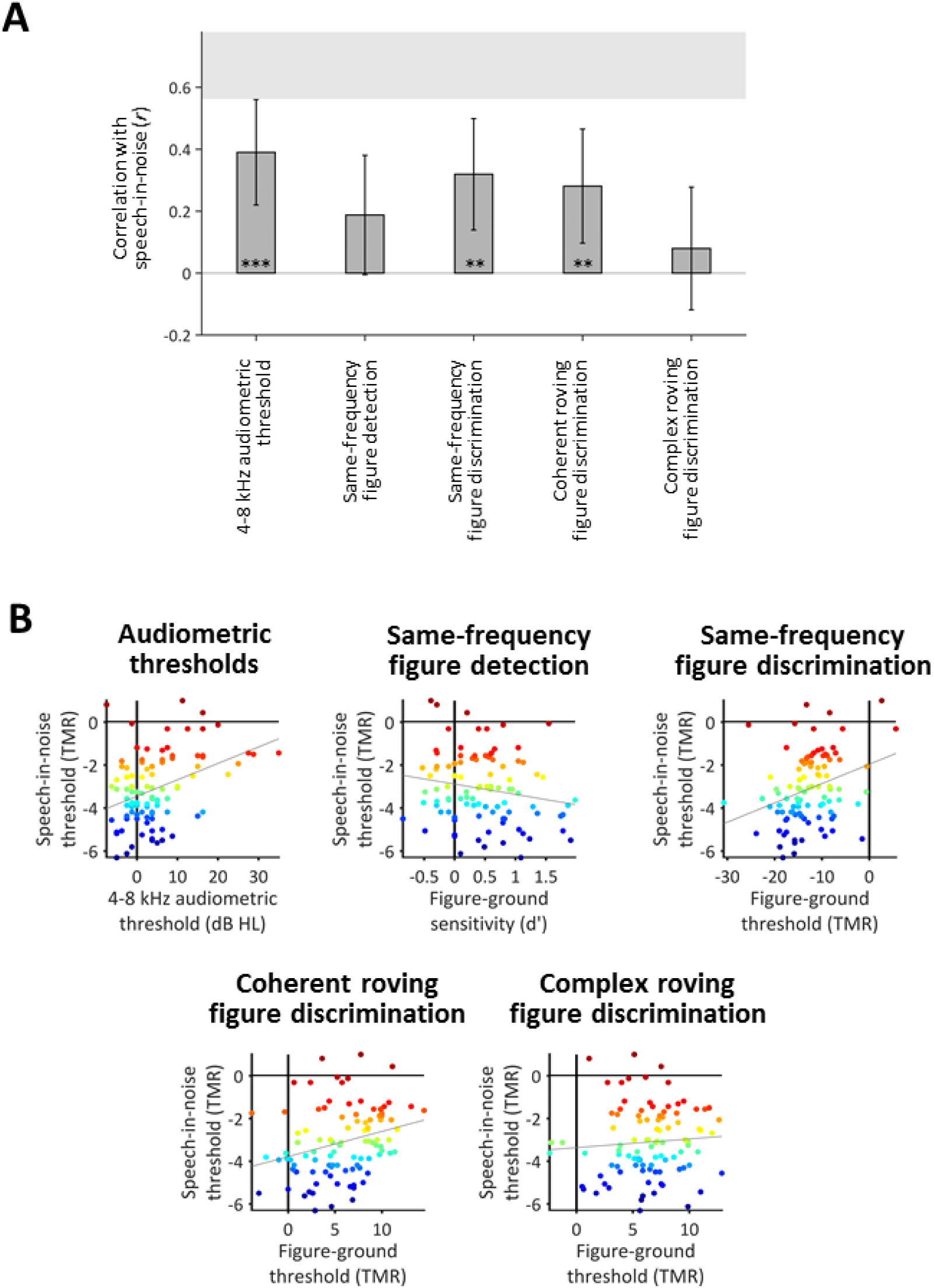
Correlations between thresholds for speech-in-babble and audiometric thresholds or thresholds for the figure-ground tasks. (A) Bar graph displaying *r*-values for Pearson’s correlations with speech-in-babble thresholds. Error bars display 95% between-subjects confidence intervals. The grey shaded box illustrates the noise ceiling, calculated as the (95% between-subjects confidence interval associated with the) correlation between two different blocks of the speech in noise task. Asterisks indicate the significance level of the correlation coefficient (* p < 0.050; ** p < 0.010; *** p < 0.001). (B) Scatter plots associated with each of the correlations displayed in Panel A. Each dot displays the results of an individual participant, which are coloured according to speech-in-noise performance (dark red = worst thresholds, dark blue = best thresholds). Solid grey lines indicate the least squares lines of best fit. [dB HL: decibels hearing level; TMR: target-to-masker ratio.] See also Supplemental Figure.

### ‘Normal’ hearing thresholds relate to speech-in-noise

Audiometric thresholds (2-frequency average at 4–8 kHz across the left and right ears) accounted for 15% of the variance in speech-in-babble thresholds (*r* = .39, *p* < .001; 95% CI = .21–.55). The correlation remained significant after excluding 8 participants who had audiometric thresholds worse than 20 dB at 4 or 8 kHz (*r* = .25, *p* = .017; 95% CI = .05–.44); excluding these participants did not lead to a significant change in the magnitude of the correlation coefficient (*z* = 1.05, *p* = 0.29), confirming that the correlation we found is driven by sub-clinical variability in audiometric thresholds.

### Different figure-ground tasks index different variance in speech-in-noise

Overall, participants achieved lower (better) thresholds in the same-frequency figure-ground discrimination task than the two roving figure-ground discrimination tasks [coherent roving figure-ground: *t*(96) = 29.07, *p* < .001, *d*_z_ = 2.95; complex roving figure-ground: *t*(96) = 29.77, *p* < .001, *d*_z_ = 3.02] (see Supplemental Figure). Most participants (N = 95) were able to perform the same-frequency task when the figure components were less intense than the background components. Whereas, for the two roving figures, most participants (Coherent roving: N = 89; Complex roving: N = 94) could only perform the task when the figure components were more intense than the background components. Thresholds did not differ significantly between the two roving figure-ground tasks [*t*(96) = 1.21, *p* = .23, *d*_z_ = 0.12].

The correlations between figure-ground thresholds and speech-in-babble thresholds are all in the expected direction (Figure 2B). Correlations with speech-in-babble were significant for the same-frequency figure-ground discrimination task (*r* = .32, *p* = .001; 95% CI = .13–.49) and the coherent roving figure-ground discrimination task (*r* = .28, *p* = .005; 95% CI = .09–.45). However, the correlation with sensitivity on the same-frequency figure-ground *detection* task (*r* = −.19, *p* = .067; 95% CI = -.37–.01) just missed the significance threshold. The correlation with the complex roving figure-ground discrimination task was not significant (*r* = .08, *p* = .44; 95% CI = −.12–.27); this is consistent with the result that thresholds in this task were unreliable between runs (i.e., within-subjects) (see Supplemental Figure).

To investigate whether performance on different figure-ground tasks explain similar (overlapping) or different variance in speech-in-babble performance, we conducted a hierarchical stepwise regression with the figure-ground tasks as predictor variables. Thresholds on the same-frequency figure-ground discrimination task accounted for 10% of the variance in speech-in-babble thresholds (*r* = .32, *p* = .001). There was a significant improvement in model fit when thresholds on the coherent roving figure-ground discrimination task were added (*r* = .39, *r*^2^ change = .05, *p* = .02); together, the two tasks explained 15% of the variance in speech-in-babble thresholds. These results demonstrate that the two figure-ground tasks explain partially independent portions of the variance—thresholds for the coherent roving figure-ground discrimination task explain an additional 5% of the variance that is not explained by thresholds for the same-frequency figure-ground discrimination task.

Possibly, the two tasks might assess the same construct, and a better fit with both tasks together is simply due to repeated sampling. To investigate this, we separated the thresholds for the two runs within each task (which were averaged in the previous analyses). We constructed models in which two runs were included from the same task (which differ in the stimuli that were presented but should assess the same construct) and models in which one run was included from each task. The idea behind these constructions is that they are equivalent in the amount of data entered into each model (always 2 runs). If the two tasks assess different constructs, the models including runs from different tasks should perform better than the models including runs from the same task. The results from these analyses are displayed in Table 1. Indeed, when one run from the same-frequency figure-ground task is entered into the model, there is no significant improvement in the amount of speech-in-babble variance explained by adding the second run (regardless of which run is entered first; upper two rows of Table 1). Whereas, adding one run of the coherent roving figure-ground task significantly improves the model fit (regardless of which run of the same-frequency task is entered first; lower two rows of Table 1). This result provides evidence that the same-frequency and coherent roving tasks assess different constructs that contribute to speech-in-noise, rather than improving model fit by sampling the same construct.

**Table 1.** Linear regression models including individual runs of the figure-ground discrimination tasks as variables. The table displays *r*-values associated with a model including variable 1 only, a model including variables 1 and 2 together, and the *r*^2^ change and *p*-values associated with adding the second variable to the model (** p < 0.01). R1: run 1; R2: run 2.

| Variable 1 | Variable 1<br>$r$-value | Variable 2 | Variable 1+2<br>$r$-value | $r^2$ change | $p$ change |
| --- | --- | --- | --- | --- | --- |
| Same-frequency (R1) | .30 | Same-frequency (R2) | .32 | .02 | .21 |
| Same-frequency (R2) | .28 | Same-frequency (R1) | .32 | .03 | .10 |
| Same-frequency (R1) | .30 | Coherent roving (R1) | .40 | .07 | .006 ** |
| Same-frequency (R2) | .28 | Coherent roving (R1) | .40 | .08 | .004 ** |

### Best predictions by combining peripheral and central measures

To examine whether the figure-ground tasks explained similar variance as audiometric thresholds, we tested models that included both audiometric thresholds and figure-ground performance. A model including the same-frequency figure-ground discrimination task and audiometric thresholds explained significantly more variance in speech-in-babble performance than audiometric thresholds alone (*r* = .45, *r*^2^ change = .05, *p* change = .016). Similarly, a model including the coherent roving figure discrimination task and audiometric thresholds explained significantly more variance than audiometric thresholds alone (*r* = .44, *r*^2^ change = .04, *p* change = .032).

When the three variables (audiometric thresholds, same-frequency figure discrimination task, and coherent roving figure discrimination task) were entered into a model together, the model explained 23% of the variance (*r* = .48). Based on our estimate of the noise in the data (defined as the correlation between thresholds measured in two separate blocks of the speech-in-babble task, reported above), the best possible variance we could hope to account for is 47%. Thus, when included together, the three tasks account for approximately half of the explainable variance in speech-in-babble performance. Although, the model including all three variables just missed the significance threshold when compared to the model including only audiometric thresholds and the same-frequency figure discrimination task (*r*^2^ change = .03, *p* change = .06).

### Figure-ground tests index age-independent deficits

Given the broad age range of participants, we examined whether task performance related to age. As expected, older age was associated with worse speech-in-noise performance (*r* = .43, *p* < 0.001; 95% CI = 0.26–0.58). Therefore, we next considered whether relationships between our tasks and speech-in-noise could be explained by age-related declines in those tasks.

Audiometric thresholds were worse in older people (*r* = .78, *p* < 0.001; 95% CI = 0.66–0.83) and the correlation between audiometric thresholds and speech-in-babble performance was non-significant after accounting for age (*r* = .10, *p* = 0.32).

Performance in the figure-ground tasks was also significantly worse in older people (Same-frequency: *r* = .23, *p* = 0.022; 95% CI = 0.03–0.41; Coherent roving: *r* = .23, *p* = 0.023; 95% CI = 0.03–0.41). However, correlations between figure-ground tasks and speech-in-babble performance were significant when accounting for age (Same-frequency: *r* = .25, *p* = 0.014; Coherent roving: *r* = .21, *p* = 0.044).

## Discussion

Our results demonstrate that people with normal hearing vary by as much as 7 dB TMR (Figure 2B) in their thresholds for understanding speech when background noise is present, and both peripheral and central processes contribute to this variability. Despite recruiting participants with audiometric thresholds that are widely considered to be ‘normal’, variability in these thresholds significantly predicted speech-in-noise performance, explaining 15% of the variance. In addition, fundamental auditory grouping processes, as assessed by our new figure-ground tasks, explained significant variance in speech-in-noise performance that was not explained by audiometric thresholds. Together, audiometric thresholds and figure-ground perception accounted for approximately half of the explainable variance in speech-in-noise performance. These results demonstrate that different people find speech-in-noise difficult for different reasons, and suggest that better predictions of real-world listening can be achieved by considering both peripheral processes and central auditory grouping processes.

### Central contributions to speech-in-noise performance

Two of the new figure-ground tests (same-frequency and coherent roving) correlated with speech-in-noise performance, and explained significant independent portions of the variance. This result suggests that (at least partially) separate processes contribute to the ability to perform the same-frequency and coherent roving tasks—and both of these processes contribute to speech-in-noise perception. That these tasks explain different portions of the variance demonstrates that they could help to tease apart different reasons that different people find it difficult to understand speech-in-noise: Some people might struggle to understand speech-in-noise due to impaired mechanisms for detecting static (same-frequency) patterns, whereas others may struggle due to impaired processes for tracking patterns that change frequency over time.

Given that neuroimaging studies show cortical contributions to both figure-ground perception (Teki et al., 2016, 2011) and to speech-in-noise perception (for a review, see Scott and McGettigan, 2013), it is highly plausible that the shared variance arises at a cortical level. That the two figure-ground tasks explain partially independent portions of the variance suggests that their explanatory power is not simply attributable to generic attention or working memory processes. Although attention and working memory may contribute to speech-in-noise perception (for example, see Akeroyd, 2008; Schoof and Rosen, 2014), we expect both of our figure-ground tasks to engage these processes to a similar extent. Consistent with this idea, same-frequency (Molloy et al., 2018; Teki et al., 2016) and roving-frequency (O’Sullivan et al., 2015) figure-ground stimuli both show neural signatures associated with figure detection during passive listening. Instead, we assume that the shared variance observed here is due to fundamental auditory grouping processes, which (at least partially) differ when a target object has a static frequency than when it changes frequency over time.

Teki et al. (2013) provide evidence that the ability to detect same-frequency figures likely relies on a temporal coherence mechanism for perceptual streaming proposed by Shamma, Elhilali, and Micheyl (2011), drawing on spectro-temporal analyses that are proposed to take place in auditory cortex (Chi et al., 2005). In this model, an ‘object’ or ‘stream’ is formed by grouping elements that are highly correlated in time across frequency channels, and these elements are separated perceptually from incoherent elements. By simulating this temporal coherence mechanism, Teki et al. (2013) show it can successfully distinguish figure-present and figure-absent trials, and provides a good fit to participants’ behavioural responses. A temporal coherence mechanism can also explain how people detect figures that change frequency over time (O’Sullivan et al., 2015), like the coherent roving stimuli used here. The three elements of the coherent roving figures change frequency in the same direction at the same rate, producing temporal coherence across frequency channels, and these would be segregated from the incoherent background of tones that have random frequencies at each time window.

Interestingly, participants obtained worse thresholds for the roving frequency figure-ground task than for the same-frequency figure-ground task: that is, the figure needed to be more intense for participants to perform the roving frequency task successfully than the same-frequency task. One plausible explanation for this finding is that both tasks require the detection of static patterns, and the roving frequency task requires additional processes for tracking frequencies over time. Yet, that the two tasks explain partially independent portions of the variance demonstrates that the processes required to perform one task are not a simple subset of the processes required to perform the other. Although a temporal coherence mechanism could be used to perform both tasks, the same-frequency and coherent roving tasks must rely on (at least partially) separate processes. For example, perhaps within-channel process are particularly important for detecting static patterns, because the frequencies of each component would fall within the same channel; whereas, processes that integrate across frequency channels might be more important for detecting roving patterns, which changed by 59–172% (interquartile range = 27%) of the median frequency of the component in this experiment. Relating these processes to speech-in-noise perception, the same-frequency figure-ground stimulus somewhat resembles the perception of vowels, which often have frequencies that remain relatively stable over time; whereas, the roving stimulus might approximate the requirement to track speech as it changes in frequency at transitions between different consonants and vowels.

The classic same-frequency figure detection task that has been used in previous studies (e.g., Molloy et al., 2018; Teki et al., 2016, 2013, 2011) did not correlate significantly with speech-in-noise perception (*p* = 0.067), but only narrowly missed the significance threshold. Although the stimuli used in the same-frequency detection task and the same-frequency discrimination task were similar, the discrimination task correlated reliably and the detection task did not. Given that the detection task used fixed stimulus parameters for every participant, whereas the discrimination task was adaptive, this result may have arisen because the adaptive (discrimination) task was marginally more sensitive to individual variability than the (detection) task with fixed parameters.

### Peripheral contributions to speech-in-noise performance

‘Normal’ variability in audiometric thresholds at 4–8 kHz explained 15% of the variance in speech-in-noise performance, suggesting that the audiogram contains useful information for predicting speech understanding in real-world listening, even when participants have no clinically detectable hearing loss. That sub-clinical variability in audiometric thresholds at these frequencies might predict speech-in-noise perception has often been overlooked: Several previous studies have assumed that sub-clinical variability in audiometric thresholds do not contribute to speech-in-noise perception and have not explored this relationship (e.g., Alvord, 1983; Kumar et al., 2012). Of those that have tested the relationship, two found a correlation (Schoof and Rosen, 2016; Yeend et al., 2017), although one of these (Yeend et al., 2017) included participants with mild hearing loss and it is possible that these participants were responsible for the observed relationship. Two others found no correlation but restricted the variability in their sample by either imposing a stringent criterion on ‘normal’ hearing less than 15 dB HL (Ruggles and Shinn-Cunningham, 2011) or considering only young participants aged 18–30 years (Oberfeld and Klöckner-Nowotny, 2016).

We infer from these results that speech-in-noise difficulties may arise from changes to the auditory periphery that are related to, but which precede, clinically relevant changes in thresholds. Although audiometric thresholds at frequencies higher than 8 kHz have been proposed to contribute to speech perception in challenging listening environments (see Pienkowski, 2016), these frequencies are not routinely measured in clinical practice. That we found correlations with 4–8 kHz audiometric thresholds suggests that speech-in-noise difficulties could be predicted based on audiometric thresholds that are already part of routine clinical assessment.

This relationship might arise because people with worse-than-average 4–8 kHz thresholds already experience some hearing loss that causes difficulties listening in challenging acoustic environments, such as when other conversations are present. Elevated high-frequency audiometric thresholds may be related directly to hair cell loss, or to cochlear synaptopathy. Regarding cell loss, even modest losses of outer hair cells could cause difficulties listening in challenging environments—for example, by degrading frequency resolution. Even people who have average thresholds between 10 and 20 dB HL have unusually low amplitude distortion product otoacoustic emissions (dpOAEs; Zhao and Stephens, 2006), which are widely considered to be related to outer hair cell dysfunction (Brownell, 1990). This finding demonstrates that even a modest loss of outer hair cells that is insufficient to produce clinically relevant shifts in audiometric thresholds has the potential to alter sound perception. On the other hand, changes in thresholds have also been suggested to accompany cochlear synaptopathy, which is an alternative mechanism that might impair speech perception. For example, Liberman et al. (2016) found that humans with a higher risk of cochlear synaptopathy, based on self-reported noise exposure and use of hearing protection, had higher audiometric thresholds at 10–16 kHz.

### Age-related changes in the auditory system

We replicated the common finding that speech-in-noise performance is worse with older age (e.g., Gordon-Salant and Cole, 2016; Humes et al., 2013). As expected, audiometric thresholds were also worse with older age, and these age-related declines in audiometric thresholds seemed to underlie their relationship with speech-in-noise performance. This is consistent with both of the possible mechanisms described above, because outer hair cells degrade with older age and post-mortem studies in humans (Makary et al., 2011; Viana et al., 2015) and animal models (Sergeyenko et al., 2013) show age-related cochlear synaptopathy.

Although performance on the figure-ground tasks became worse with older age, the relationship between figure-ground and speech-in-noise performance remained significant after accounting for age. This finding suggests that age-independent variability in figure-ground perception contributes to speech-in-noise performance. That is, even among people of the same age, figure-ground perception would be expected to predict speech-in-noise performance. An interesting question for future research would be to explore whether the shared age-independent variance is because some people are inherently worse at figure-ground and speech-in-noise perception than others, or whether these same people begin with average abilities, but experience early-onset age-related declines in performance.

### Clinical applications

A sizeable proportion of patients who visit audiology clinics report difficulties hearing in noisy places, despite having normal audiometric thresholds and no apparent cognitive disorders (Cooper and Gates, 1991; Hind et al., 2011; Kumar et al., 2007)—and currently there is no satisfactory explanation for these deficits. Although the current experiment sampled from the normal population, our figure-ground tests might be useful for assessing possible central grouping deficits in these patients. These patients are sometimes diagnosed with ‘auditory processing disorder’ (APD), despite little understanding of the cause of their difficulties or ways in which we can help these patients. Nevertheless, children with APD have speech-in-noise perception that appears to be at the lower end of the normal range (Ferguson et al., 2011). If these patients also perform poorly on figure-ground tests, then future research might focus on testing strategies to improve fundamental grouping processes.

The figure-ground tasks we developed were quick to run (∼10 minutes each), making them feasible to add to standard clinical procedures alongside the pure-tone audiogram. These tests may help clinicians gain a better understanding of the types of deficits these patients face, as well as helping to predict real world listening beyond the audiogram. Furthermore, performance in these tasks is independent of linguistic ability, unlike standard speech-in-noise tests. They would therefore be appropriate for patients who are non-native speakers of the country’s language (which is important given that speech-in-noise perception is worse in a listener’s second than native language; e.g. Mayo et al., 1997), and for children who do not have adult-level language skills. Given that clinical interventions, such as hearing aids and cochlear implants, typically require a period of acclimatisation before patients are able to successfully recognise speech, these tests which use simple pure tones may be useful for predicting real-world listening in the early stages following clinical intervention.

One step towards clinical application would be to improve the reliability of these measures (see Supplemental Figure), which might allow them to explain even greater variability in speech-in-noise performance than reported here. Between-run variability in figure-ground thresholds could be due to stimulus-specific factors, such as frequency separation. We suspect the reason that the complex roving figure-ground task did not correlate with speech-in-noise performance was because it was unreliable within participants (between run correlation: *r* = −0.02); a possible alternative explanation that it was more difficult than the other tasks was not supported by the data. The complex roving task may vary more than the other figure-ground tasks because it contains additional variability related to the second and third formants of a spoken sentence, and these components (including the frequency changes within each component and their relationship to the frequency changes in the first formant) might impact the extent to which the figure can be extracted; in contrast, variability in the extent to which the first formant changes frequency is present in both the coherent and complex roving figures.

## Conclusions

Overall, our results demonstrate that speech-in-noise difficulties can occur for a variety of reasons, which are attributable to impairments at different stages of the auditory pathway. We show that successful speech-in-noise perception relies on audiometric thresholds at the lower end of the normally hearing range, which likely reflect differences at the auditory periphery. Our results also reveal that fundamental grouping processes, occurring centrally, are required for successful speech-in-noise perception. We introduce new figure-ground tasks that help to assess the grouping of static acoustic patterns, and the ability to track acoustic sources that change in frequency over time— interestingly, both of these processes appear to be important for speech-in-noise perception. These findings highlight that speech-in-noise difficulties are not a unitary phenomenon, rather suggesting that we require different tests to explain why different people struggle to understand speech when other sounds are present. Assessing both peripheral (audiometric thresholds) and central (grouping) processes can help to characterise speech-in-noise deficits.

## Materials and Methods

### Subjects

103 participants completed the experiment. We measured their pure-tone audiometric thresholds at octave frequencies between 0.25 and 8 kHz in accordance with BS EN ISO 8253-1 (British Society of Audiology, 2004). We excluded 6 participants who had pure-tone thresholds that would be classified as mild hearing loss (6-frequency average ≥ 20 dB HL in either ear). We analysed the data from 97 participants, which we determined would be sufficient to detect significant correlations of *r*^2^ ≥ 0.12 with .8 power (Faul, Erdfelder, Lang, & Buchner, 2007). The 97 participants (40 male) were 18–60 years old (median = 24 years; interquartile range = 11). The study was approved by the University College London Research Ethics Committee.

### Experimental Procedures

The experiment was conducted in a sound-attenuating booth. Participants sat in a comfortable chair facing an LCD visual display unit (Dell Inc.). Acoustic stimuli were presented through an external sound card (ESI Maya 22 USB; ESI Audiotechnik GmbH, Leonberg) connected to circumaural headphones (Sennheiser HD 380 Pro; Sennheiser electronic GmbH & Co. KG) at 75 dB A.

Participants first performed a short (< 5 minute) block to familiarise them with the figure-ground stimuli. During the familiarisation block, they heard the figure and ground parts individually and together, with and without a gap in the figure.

Next, participants completed 6 tasks (Figure 1): four figure-ground tasks and two blocks of a speech-in-babble task. All tasks were presented in separate blocks and their order was counterbalanced across participants. Immediately before each task began, participants completed 5 practice trials with feedback. No feedback was provided during the main part of each task.

One of the figure-ground tasks was based on a detection task developed by Teki et al. (2011), in which the stimuli consisted of 40 50-ms chords with 0 ms inter-chord interval. Each chord contained multiple pure tones that were gated by a 10-ms raised-cosine ramp. The background comprised 5– 15 pure tones at each time window, whose frequencies were selected randomly from a logarithmic scale between 179 and 7246 Hz (1/24^th^ octave separation). The background lasted 40 chords (2000 ms). For the figure, we used a coherence level of 3 and a duration of 6. The frequencies of the 3 figure components were also selected randomly, but with an additional requirement that the 3 figure frequencies were separated by more than one equivalent rectangular bandwidth (ERB). The frequencies of the figure were the same at adjacent chords. The figure lasted 6 chords (300 ms) and started on chord 15–20 of the stimulus. For half of stimuli, there was no figure in the stimulus; to ensure that figure-present and figure-absent stimuli had the same number of elements (and therefore the same amplitude), figure-absent stimuli contained an additional 3 components of random frequencies, which had the same onset and duration as the figures in figure-present stimuli. Participants’ task was to decide whether the figure was present or absent on each trial. Each participant completed 50 trials, with an inter-trial interval between .8 and 1.2 seconds.

In three of the figure-ground tasks, participants completed a two-interval two-alternative forced choice discrimination task. On each trial, participants heard two figure-ground stimuli sequentially, with an inter-stimulus interval of 400 ms. Both stimuli contained a figure that lasted on average 42 chords (2100 ms) and a background that lasted exactly 3500 ms (70 chords). For one stimulus, 6 chords (lasting 300 ms) were omitted from the figure. For the other stimulus, the same number of components (3) were omitted from the background (6 chords; 300 ms). Participants’ task was to decide which of the two stimuli (first or second interval) had a ‘gap’ in the figure. In the “same-frequency” task, the figure lasted exactly 42 chords (2100 ms) and the 3 figure components were the same frequencies at adjacent chords, similar to the figure-ground detection task. In the “complex roving” task, the 3 figure components were based on the first three formants of the sentences used in the speech-in-noise tasks. We extracted the formants using Praat (http://www.fon.hum.uva.nl/praat/), and averaged the frequencies of the formants in 50-ms time bins; we then generated 50-ms pure tones at those frequencies. In this task, the figure lasted for the same duration as the extracted formants (34–50 chords; median = 42 chords; interquartile range = 4). In the “coherent roving” task, the 3 figure components were multiples of the first formant frequencies: the first component was equal to the first formant frequency, the second component was the first component multiplied by the average difference between the first and second formants in the sentence, and the third component was the second component multiplied by the average difference between the second and third formants. In all three tasks, we varied the target-to-masker ratio (TMR) between the figure and ground in a 1-up 1-down adaptive procedure (Cornsweet, 1962) to estimate the 50% threshold. Each run started at a TMR of 6 dB. The step size started at 2 dB and decreased to .5 dB after 3 reversals. For each task, we adapted the TMR in two separate but interleaved runs, which were identical, except that different stimuli were presented. Each run terminated after 10 reversals.

Participants completed two blocks of the speech-in-noise task, which each contained two interleaved runs; these were identical, except different sentences were presented as targets. Sentences were from the English version of the Oldenburg matrix set (HörTech, 2014) and were recorded by a male native-English speaker with a British accent. The sentences are of the form “<Name> <verb> <number> <adjective> <noun>” and contain 10 options for each word (see Figure 1). An example is “Cathy brought four large chairs”. The sentences were presented simultaneously with 16-talker babble, which began 500 ms before the sentence began, ended 500 ms after the sentence ended, and was gated by a 10-ms raised-cosine ramp. A different segment of the noise was presented on each trial. Participants’ task was to report the 5 words from the sentence (in any order), by clicking words from a list on the screen. The sentence was classified as correct if all 5 words were reported correctly. We adapted the TMR between the sentence and babble in a 1-up 1-down adaptive procedure, similar to the figure-ground discrimination tasks. The TMR began at 0 dB and the step size started at 2 dB, which decreased to .5 dB after 3 reversals. Each run terminated after 10 reversals.

### Analyses

For the figure-ground detection task, we calculated sensitivity (d′; Green and Swets, 1966) across all 50 trials.

For the adaptive tasks (speech-in-babble and figure-ground discrimination tasks), we calculated thresholds as the median of the last 6 reversals in each run. For the main analyses, the thresholds from the two interleaved runs within each block were averaged.

To isolate the contributions of different tasks to speech-in-noise, we used a hierarchical linear regression with the stepwise method. All correlations are Pearson’s correlation coefficients, reported without correction, given that the conclusions of the paper are based on the results of the regression analyses rather than the *p*-values associated with the correlations.

To compare average thresholds between different figure-ground discrimination tasks, we used paired-samples *t*-tests.

## Additional Information

### Funding

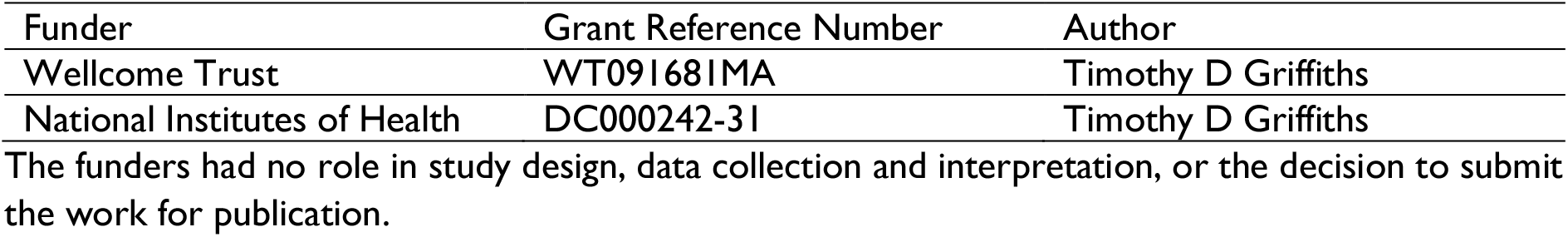

### Author Contributions

EH: Conception and design, Acquisition of data, Analysis and interpretation of data, Drafting or revising the article; TDG: Conception and design, Analysis and interpretation of data, Drafting or revising the article.

### Ethics

Human subjects: 103 volunteers gave informed written consent, in accordance with institutional guidelines. This study was approved by the University College London Research Ethics Committee.

