## Supplementary figures and images for "‘Normal’ hearing thresholds and fundamental auditory grouping processes predict difficulties with speech-in-noise perception"

### Supplemental Figure

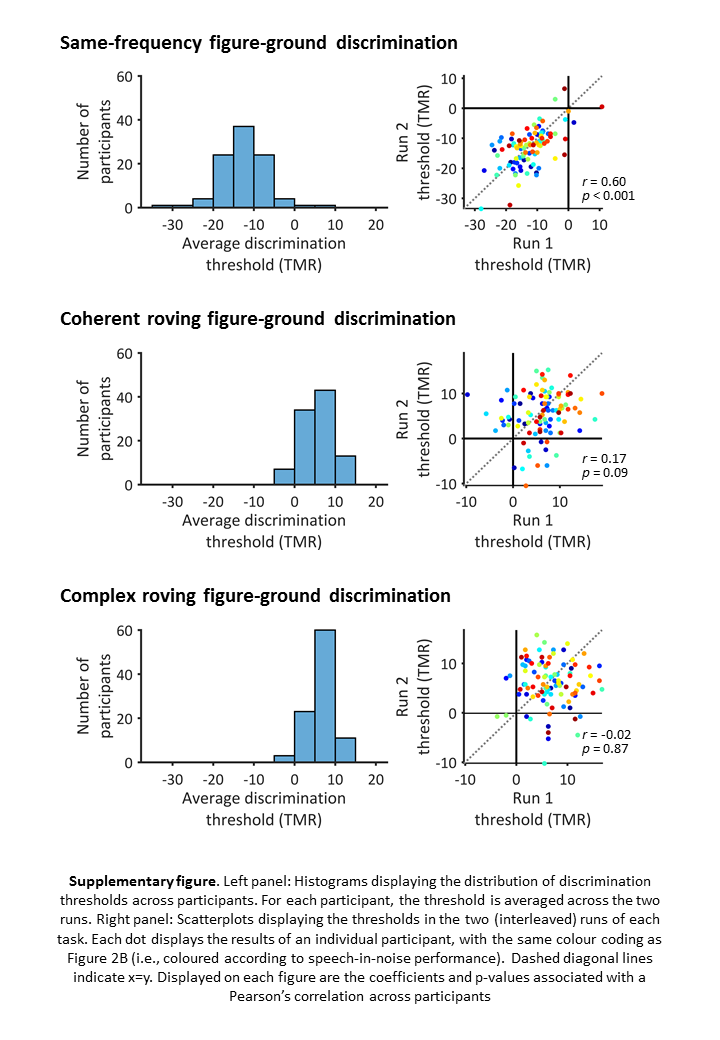
